# Introducing undergraduate students to reverse genetics in the laboratory: a case study of a project-oriented learning (POL) system in Taishan Medical University, China

**DOI:** 10.1101/2020.05.21.108068

**Authors:** Lijuan Yu, Henglong Zhang, Jing Zhai

## Abstract

Biochemistry and molecular biology have become increasingly reliant on experimental experience, because they are difficult subjects to understand from a theoretical approach. The Innovation and Entrepreneurship Training Program was established by the China Ministry of Education to foster the scientific interests and experiences of undergraduate students. Here, we reported on an innovative project-oriented learning (POL) system directed by Taishan Medical University (Named Shandong First Medical University & Shandong Academy of Sciences since February 2019) China, which fused innovative project with biochemical course concepts. This study takes reverse genetics, which is considered to be difficult to understand through course-only learning, as an example. To help undergraduate students understand the principle of reverse genetics, the project involved producing hepatitis C virus *in vitro*. Furthermore, to give them a comprehensive scientific experience based on their knowledge from textbook, extensive mentorship was provided through the project, including theoretical guidance and technical support for experiments. Assessment showed a comprehensive improvement after participating in these programs. In conclusion, this POL system was a good model that enabled undergraduates to learn more during courses. It also provided a chance to experience hands-on sciences, which might motivate them to pursue a career in science.

---

Following China’s economic development and industrial transformation, the innovation-driven approach has gradually become a national development strategy. The cultivation of scientific and technological innovative talents is the key force to promote this process. Compared to western countries, China has alarge population of undergraduates, but its innovation atmosphere is relatively low. Engaging undergraduates in the practices of sciences is seen as a critical factor for preparing the next generation scientists. In terms of biomedical education, Chinese government has recently provided increasing support for scientific training program for undergraduates. Over 50,000 grants written by undergraduate students were approved in the period 2012-2016 by the Student’s Platform for Innovation and Entrepreneurship Training Program, which was established by the China Ministry of Education (CME) to foster the scientific interests of undergraduates. Approved students can conduct their proposed projects under the supervision of mentors. This approach was a good way to involve students and give them primary scientific experience as well as motivate them to be more active in academic research. It has been reported that the students involved in the program are better trained as research scientists and are more likely to pursue graduate studies/careers in the sciences, compared to non-participating students [1-3]. This group of highly motivated and trained undergraduates might make significant contributions to Chinese scientific developments in the future.

Biochemistry and Molecular biology have been increasingly reliant on experimental experience, because they are difficult to understand from a theoretical approach. However, many new concepts were difficult to incorporate into the undergraduate experimental course setting, which has created a lack of training in molecular biology, which affects undergraduates’ future career development in medical school candidates or as biologists. Many reports have called for a “revolution” in undergraduate molecular biology education, in which experimental skills receive a greater emphasis.

Usually, teachers explain concepts during class and might combine these lectures with an animation in order to help students understand a subject more clearly. However, students might still be confused. The Experimental opportunities provided by government-approved innovative project-oriented learning (POL) could provide students with chances to perform the experiments on their own. This would have a more direct impression. Here, we report on “reverse genetics” which represents the undergraduate training program, to show how this project progressed in a medical university and how it benefited the teaching of biochemistry and molecular biology.

Every year, about 700 students majoring in medical science enroll into Taishan Medical University (TSMU). In the biochemistry course, the teachers address several topics related biochemistry theory, such as “reverse genetics,” “tumorigenesis,” “metabolic disorder,” “protein structures,” “gene recombination,” and “enzymatic catalyzation.”Among the students, approximately 20% can obtain approval for the innovative project with suitable topics. These students are divided into several groups. They then choose their instructors based on their course topics and project. The innovation team itself is composed of individual student and instructors. And the instructor and the students (usually one instructor for1-5 students in a team)perform experiments together as a team in the instructor’s laboratory.In this study, we selected “reverse genetics”as an example to show how the innovative POL system works. The project focuses on the *in vitro* production of hepatitis C virus (HCV), a positive-strand Ribonuclease Acid (RNA) virus that belongs to the Hepacivirus genus in the Flaviviridae family [4-5]. The laboratory culture strain (named JFH1-B/WT) and the related culture system form an appropriate model to introduce undergraduates to reverse genetics. The complete procedure used to produce HCV *in vitro* consists of four steps: plasmid Deoxyribonucleic Acid (DNA) linearization, DNA transcription, RNA transfection to the cell, and virus imaging (Fig. 1). The information provided was kept as straightforward as possible to accommodate the basic level of students’ scientific backgrounds. The following are examples of background information provided to the students.

**Fig. 1.**
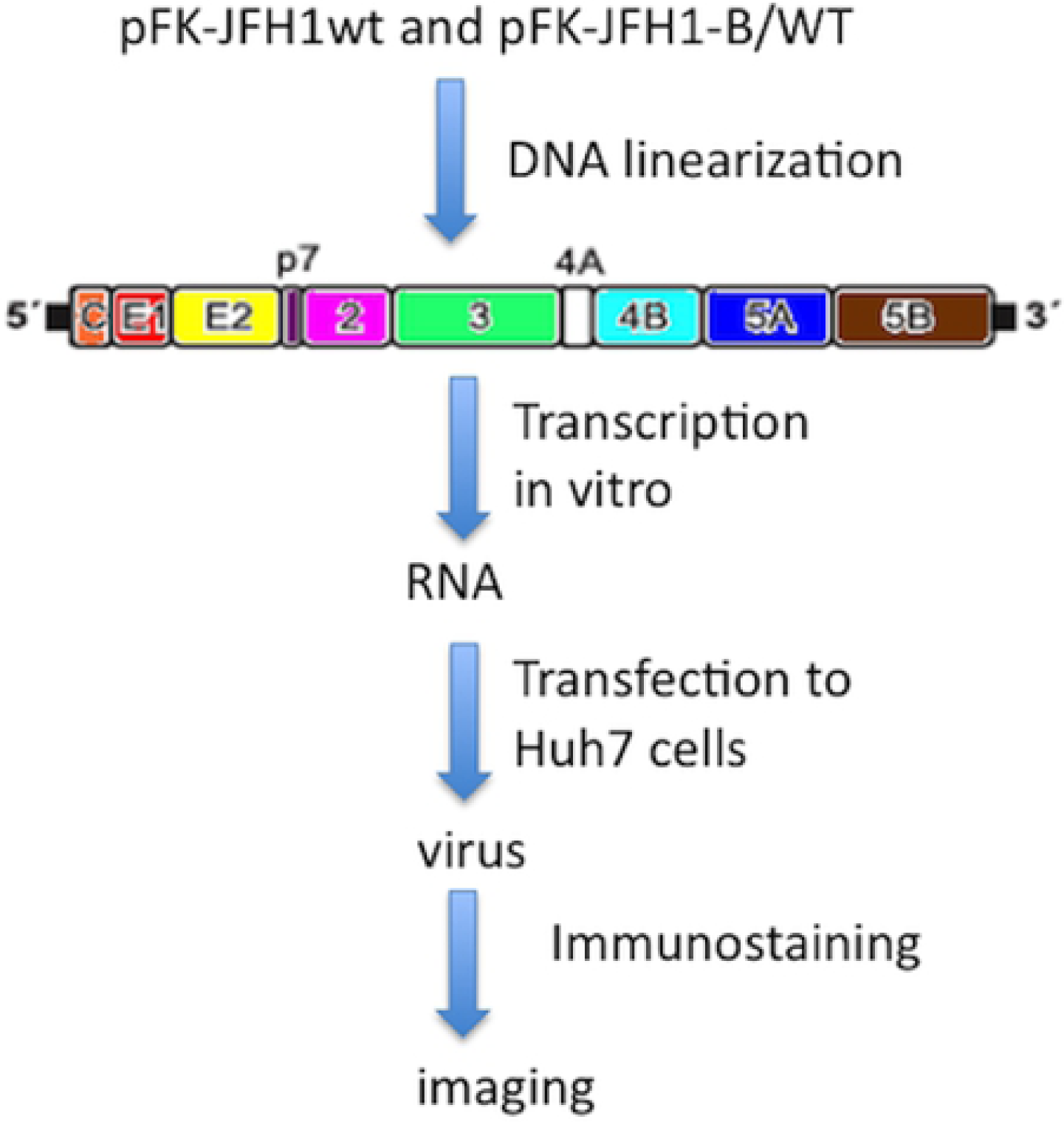
General experiments outline of the project. JFHl-WT and JFHl-B/WT plasmid were digested to generate linearized DNA, which would be used as template for synthesis RNA in vitro. The RNAs then were transfected into Huh7 cells to produce viruses, which was furthure imagined by immunostaining.

“Reverse genetics is an approach to investigate the function of a specific gene by analyzing the phenotypic effects of specific engineered gene sequences. While forward genetics seeks to find the genetic basis of a phenotype or trait, reverse genetics seeks to find what phenotypes arise as a result of particular genetic sequences. To learn the influence that sequence has on phenotype or to discover its biological function, researchers can engineer a change or disruption in the DNA by mutagenesis. The researchers can look for the effect of such alterations in the whole organism.

“Immortal carcinoma cell culture is one of the major in vitro model systems used for studying cellular and molecular biology. One use of cultured cells is to act as a host system for viruses. Compared to animal models used in virology, the main advantages of the cell culture system are low expense, easier monitoring and better reproducibility of results obtained”.

“JFH1 (Japanese fulminant hepatitis) is a widely used laboratory virus strain that could replicate efficiently in Huh7 (liver carcinoma cell line)[6-9].The JFH1-B/WT is a variant derived from JFH1. Compared to JFH1, several mutations are detected by sequencing analysis in the genome of JFH1-B/WT [10], which may cause the phenotype change according to the basic ‘central dogma’.”

## Experiment implementation

### Participants and safety issues

A group consisting of one student and one mentor was selected to study the POL. The first student was a 3^rd^ year undergraduate of the TSMU who received a grant (Student’s Platform for Innovation and Entrepreneurship Training Program, China Ministry of Education. #201510439112) from the CME in 2015. He was quite interested in molecular biology and totally involved in most of the experiments in the project. Before performing experiments in the laboratory, he received safety training from the laboratory manager of the university and passed a test. The mentorwas an assistant professor responsible for directing the student to conduct the experiments properly. The experiments related to live pathogens (transfection and virus staining) were performed by the mentor, as the 3^rd^ year undergraduate student did not have experience in performing experiments in a Biosafety2+ Laboratory. All the experiments were performed in the laboratory for scientific research of biochemistry department. This study was approved by the Medical Ethics Committee of Shandong First Medical University & Shandong Academy of Medical Sciences and all participants informed consent when the data collected.

### Cell culture

The human hepatoma cell line Huh7 cells [11-12] were cultured in Dulbecco’s modified Eagle’s medium (DMEM) (Thermo Scientific, Hudson, NH) supplemented with 10% fetal bovine serum (Invitrogen Corporation/Gibco, CA), 100 U/ml penicillin and streptomycin 100 μg/ml (American Type Culture Collection, Manassas, VA), and non-essential amino acids (Invitrogen Corporation/Gibco, CA). All cells were passaged at 80-90% confluence for long-term maintenance in culture.

### Linearization of plasmid DNA

10 μg JFH1 or JFH1-B/WT plasmid [10] was linearized with MluI (NEB, CA) for 3 hrs at 37°C. The complete linearization of DNA was confirmed by gel electrophoresis (Fig. 1), and linearized DNAs were purified with phenol/chloroform followed by ethanol precipitation. The integrity and concentration of recovered linearized DNA was quantified at OD.280 with spectrophotometerND-1000 (NanoDrop Technologies, USA), and the linearized DNA was used as the template for RNA synthesis.

### RNA synthesis *in vitro*

The RNA synthesis was performed as the manufacturer’s description (Applied Biosystems/Ambion, TX). In brief, the transcription reaction was performed at room temperature, it was mixed thoroughly and incubated at 37°C for 4 hrs; later, 1 μl TURBO DNase was added and incubated for 15 min at 37°C to remove the template DNA. The RNAs synthesized *in vitro* were purified using an RNA extraction kit (Omega, CA).

### Virus production

The Huh7 cells, which were at an 80% confluency, were transfected using Lipofectamine 2000 (Invitrogen, CA). In brief, Liposome-mediated transfection was performed with Lipofectamine 2000 (Invitrogen, CA) at an RNA lipofectamine ratio of 1:2 by using 4 g of JFH1-B/WT RNA in cell suspensions containing 106 cells. Cells were then plated in DMEM with 20% FCS for overnight incubation [11]. The culture supernatants were harvested 72hrs post-transfection and used to infect naive Huh7 cells.

### Indirect fluorescence immunostaining of infected cells

Tens of thousands of Huh7 cells/well were seeded in96-well plates. 24 hrs later, the cells were infected with 100μl of serially diluted culturing supernatants (collected after 72 hrsof transfection). 72hrs later, the infected cells were rinsed once with PBS, fixed with 10%formaldehyde, and permeabilized with 0.5% TritonX-100/PBS. Immunostaining was performed with mouse anti-HCV core antibody (Clone Hyb-K0811B, Cosmo Bio, Tokyo, Japan) (1:200) for 1 h at room temperature. The primary antibody was detected by a 2^nd^ antibody labeling with anti-mouse-Alexa Fluor 488 (Santa Cruz, CA).

## Results

Reverse genetics isa concept described in molecular biology.Without any scientific and laboratory background, undergraduates would be unlikely to understand it properly, even if they were to get a high score on the final tests. Therefore, the POL was introduced.

From 2012 to2016, more than 400 grants written by undergraduates were approved at TSMU. Systematical analysis showed that students with experience in scientific work have a higher chance of writing a grant by themselves and getting approved than other students. We reported a combined-project case study, supported by grants, featuring both the mentor and undergraduate. Both were actively involved, as they were passionate about the scientific field of molecular biology. In addition, high-quality mentorship could result in more efficient project achievements [12].

By participating in this project, students learnt about gene mutations and their corresponding effects step by step. First, the experimental outline was designed, as shown in Fig. 1. The DNA plasmids JFH1-WT and JFH1-B/WT were digested by restriction endonuclease MluI, and the complete digestion efficiency was checked by agarose gel electrophoresis, as shown in Fig. 2.Next, the products were used as the template to synthesize RNA *in vitro*, and the RNA products were confirmed by gel electrophoresis, as shown in Fig. 3.The Huh7 cells were then transfected by RNA with Lipofectamine 2000. Seventy-two hours later, the culture supernatants were collected and used to infect naïve Huh7 cells for viral imaging. As shown in Fig. 4, both the produced live HCV numbers and morphology were changed by comparing the WT and B/WT virus images. This indicated that the mutations had a genetic influence on the viral phenotype, at least for the morphology and replication efficiency. This phenomenon was an adjustable example that could represent the laboratory practice of the described reverse genetics concept.

**Fig. 2.**
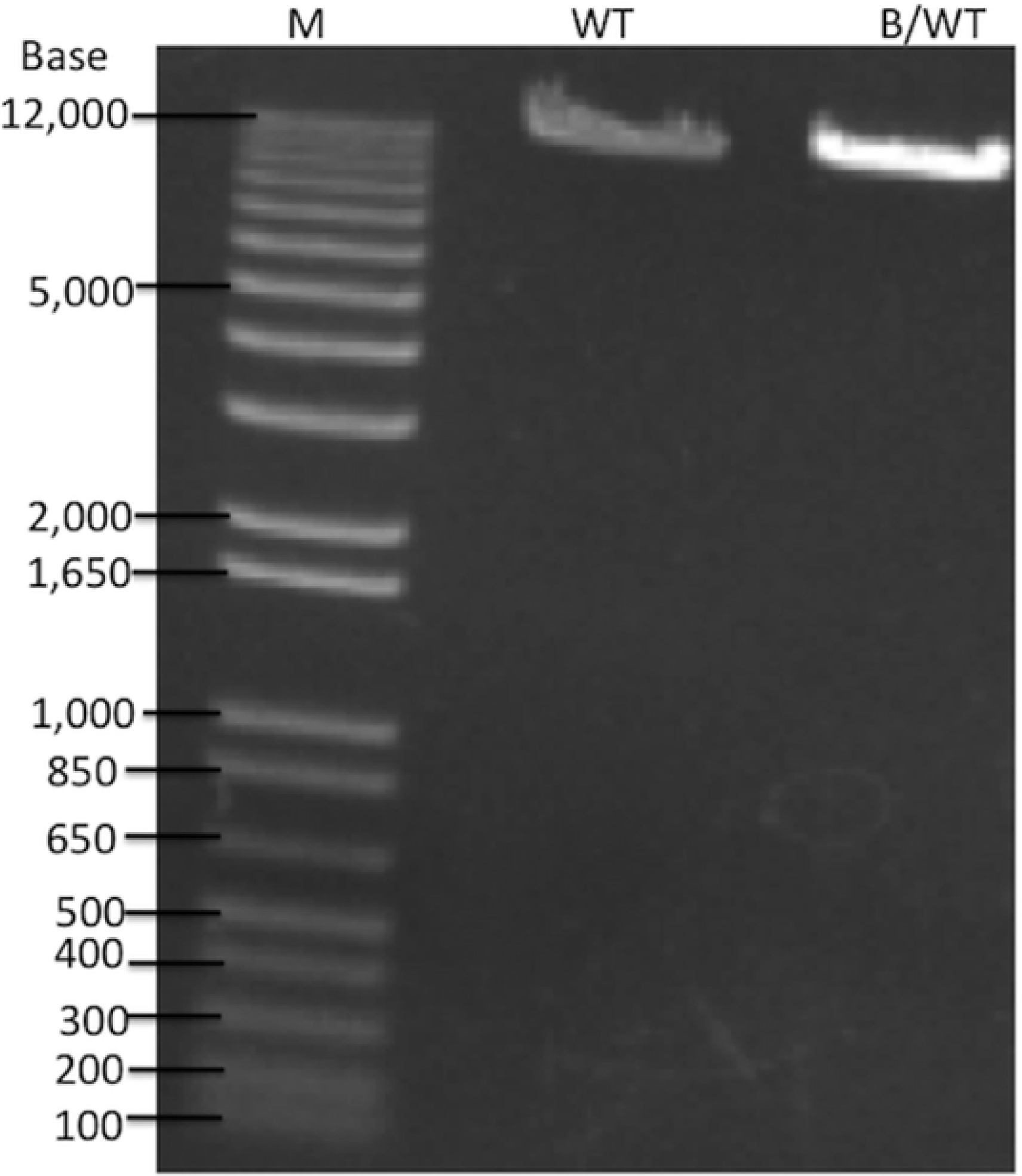
The agarose gel electrophoresis of linearized DNA. The pFK-JFHl-WT and pFK-JFHl-B/WT were digested by MluI for 3hrs at 37 °C, then the 5ul digested products were applied to 1%agarose gel for electrophoresis. M represents DNA ladder (Thermofisher 1Kb plus DNA Marker), WT and B/WT represent JFHl-WT and JFHl-B/WT individually.

**Fig. 3.**
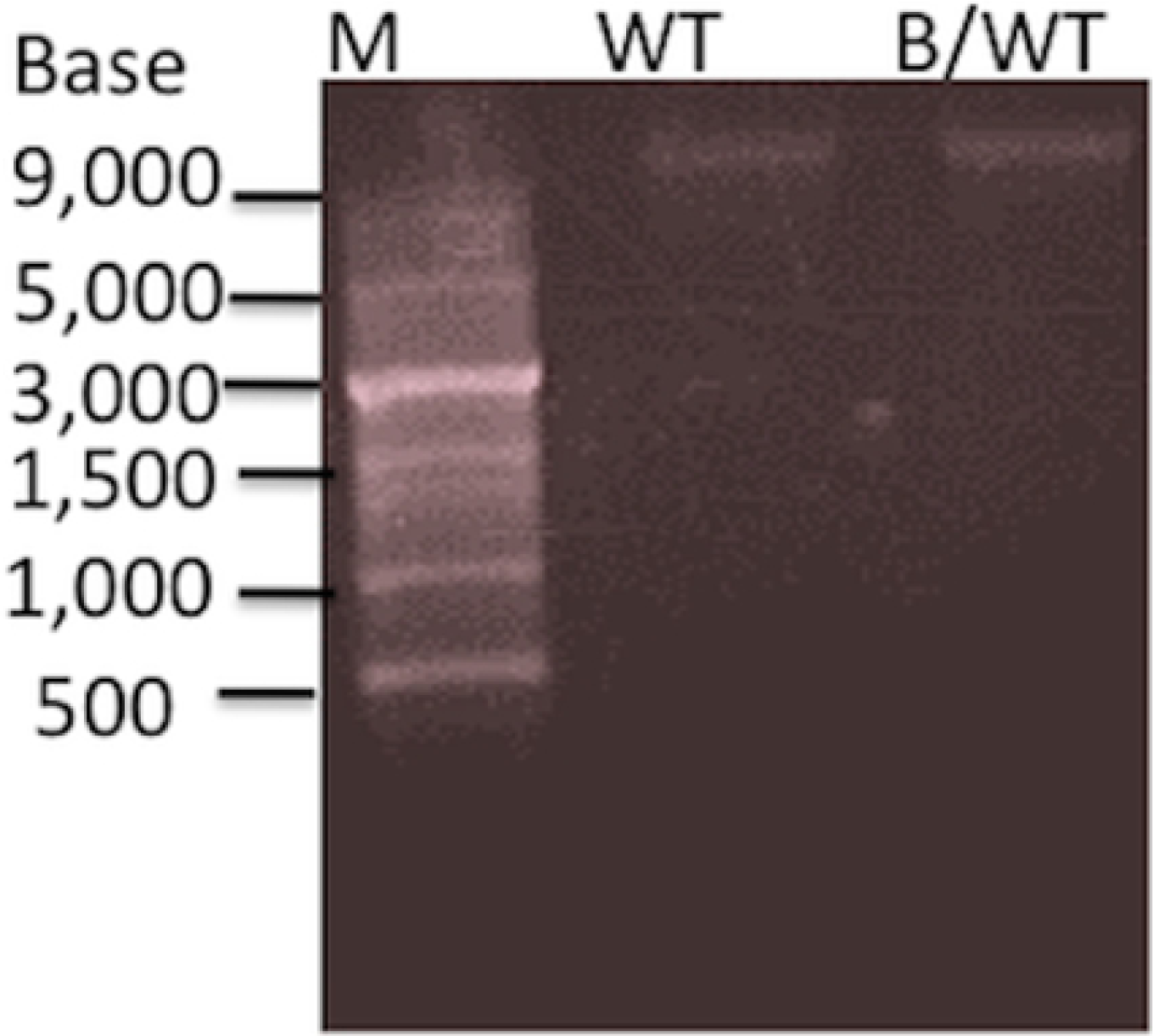
The agarose gel electrophoresis of RNA synthesized in vitro. The RNAs synthesized mixed with loading buffer at ratio 1:1 were denatured at 95 °C for 5 min, then immediately placed on ice for 2min. Then the RNAs were loaded in 1.5% gel for electrophoresis. M represents the RNA ladder (Takara), WT and B/WT represent the JFH1WT and JFH1-B/WT individually.

**Fig. 4.**
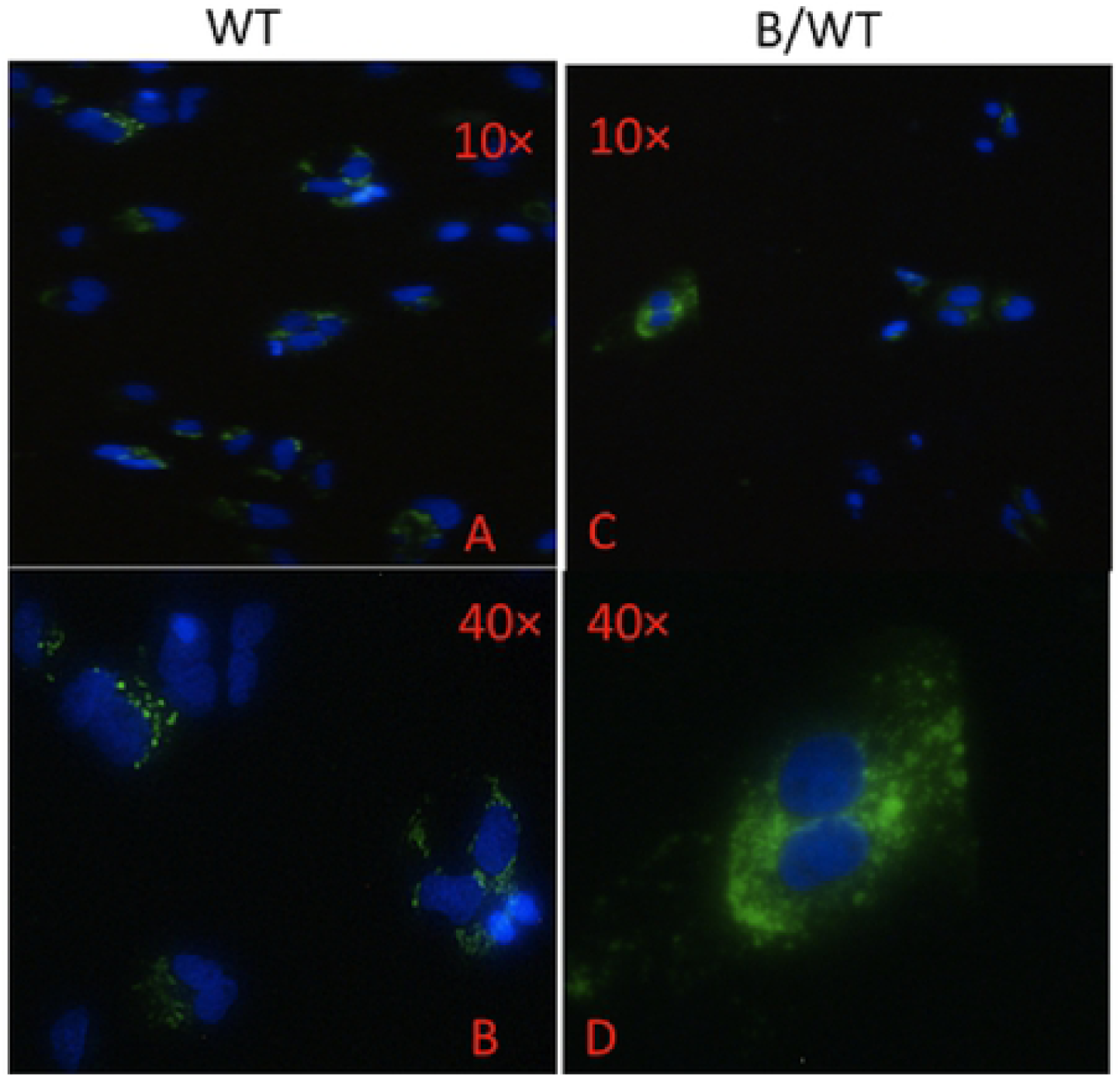
Immunofluorescence of infected Huh7 cells. The Huh7 cells splitted 24hrs before were infected with virus containing supernatant, 72hrs post infection infected cells were fixed with acetone for indirect immunofluorescence staining. The virus was visualized in green by anti-HCV core primary antibody followed by labeling with anti-rabbit-Alexa Fluor 488 secondary antibody. The nucleus was stained by DAPI in blue. The panel A and panel Cwere images of cells infected by WT and taken photo in 10 and 40 times microscopy individually, the panel B and panel D were images of cells infected by WT and taken photo in 10 and 40 times microscopy individually.

In past years, the course-based undergraduate research experiences (CUREs) have emerged as a viable mechanism to increase novices’ development of scientific reasoning and processing skills in the disciplines of science, engineering, technology and mathematics [13]. CURE usually enrolls large-scale undergraduates in a short-term project [14-15]. In our innovative POL model, the individual group would perform their individual experiments in a small scale, long-term project, and the faculty could provide one-on-one supervision of undergraduate researchers. In the case study of “reverse genetics” grant, we incorporated the concept of “reverse genetics” into innovative project based on biochemistry class. Resources were used and experiments performed in the instructors’ laboratory, rather than undergraduate labs. Compared to traditional experiment courses, in which students follow specified study designs and protocols to demonstrate well-known and understood principles or phenomena,this model had an advantage in that it let students participate in autonomous exploration that was both iterative and discovery-based in nature, at an early stage in their study (second year in university).This approach will likely encourage their interests to science, even providing new horizon to biochemistry course-related concepts. This constitutes a harmonious fusion between scientific research and medical course education. Furthermore, students involved in different projects would tackle different detailed problem, allowing them to discuss these problems under the direction of their instructor until they found the reasonable answers. Thus, the comprehensive capability of students in different specific fields was enhanced. As CURE has focused extensively on student outcomes, recent efforts have tremendously increased official regulation and formal training in the responsible conduct of research (RCR)[16]. The overarching goal of these initiatives is to ensure that a majority of researchers receive some form of RCR instruction in their careers. In terms of POL model, young undergraduates have obtained relevant experiences (e.g., mentoring by a senior scientist) since they joined in the team. In addition, the mentorship required a much higher level, as it involved long-term participation. Regarding individual teams, this model could also be directly targeted to address an individual’s needs and interests, and was quite adaptable to reproducibility since it was small in scale.

## Assessment and Conclusion

To evaluate the outcomes of POL, we introduced the Undergraduate Student Self-Assessment (a questionnaire) in a relative large scale. In total, 112 students from 34 innovation groups participated in and completed the questionnaires, as shown in Table 1.The questionnaires contained 10 self-assessment items, which were used to indicate undergraduates’ self-assessed gains (defined as good or moderate or poor). These results showed that100% students thought that the model could improve the communication skills, 99% thought it could increase their collaboration skills, 91% thought it could enhance self-efficiency, 91% thought it could enhance their science identity, 100% thought it could increase access to mentorship and motivation, 99% thought it could increase resilience, 95% thought it could increase their ability to navigate uncertainty and increase their socioemotional support, 100% students thought it could enhance their ability in understanding concepts of biochemistry and molecular biology course. Compared to most studies on CUREs, the assessment of POL involved different groups and was relatively more reliable due to the crosstalk [17].According to the feedback provided by this questionnaire, all the other items were enhanced by more than 95%except for self-efficiency and science identity, which increased by 91%. Students’ comprehensive literacy also largely improved. The POL used the theories that the students learned and developed to guide the design and implementation of studies. The funds used in POL were supported by Chinese government and were efficiently utilized. The model was therefore quite effective in cultivating undergraduates’ capacity to become more sophisticated science practitioners. Regarding future studies, several interesting aspects of POL should be carefully analyzed. For example, currently there is limited time to implement big changes to the curriculum. Faculties would prefer to adopt readily available impressive model systems (such as reverse genetics) to plug into the POL like most CUREs did, but these systems lacked the collaboration necessary to promote long-term authentic research evolution [18]. In terms of government-approved POL, China Ministration of Education may be well positioned to provide the variety and depth of projects necessary to sustain POL.

## Funds

This work is funded by the natural science foundation of Shandong province (ZR2014HQ013),the scientific development project of Shandong Health administration(2017ws157), the Shandong Traditional Chinese Medicine Administration (2015-257), and the Tai’an city (2015-NS2093), undergraduate innovative grant (201510439112) from Ministration of Education, China.

